# Bayesian Inference of Joint Coalescence Times for Sampled Sequences

**DOI:** 10.1101/2021.07.23.453461

**Authors:** Helmut Simon, Gavin Huttley

## Abstract

The site frequency spectrum (SFS) is a commonly used statistic to summarize genetic variation in a sample of genomic sequences from a population. Such a genomic sample is associated with an imputed genealogical history with attributes such as branch lengths, coalescence times and the time to the most recent common ancestor (TMRCA) as well as topological and combinatorial properties. We present a Bayesian model for sampling from the joint posterior distribution of coalescence times conditional on the SFS associated with a sample of sequences in the absence of selection. In this model, the combinatorial properties of a genealogy, which is represented as a coalescent tree, are expressed as matrices. This facilitates the calculation of likelihoods and the effective sampling of the entire space of tree structures according to the Equal Rates Markov (or Yule-type) measure. Unlike previous methods, assumptions as to the type of stochastic process that generated the genealogical tree are not required. Novel approaches to defining both uninformative and informative prior distributions are employed. The uncertainty in inference due to the stochastic nature of mutation and the unknown tree structure is expressed by the shape of the posterior distributions. The method is implemented using the general purpose Markov Chain Monte Carlo software PyMC3. From the sampled posterior distribution of coalescence times, one can also infer related quantities such as the number of ancestors of a sample at a given time in the past (ancestral distribution) and the probability of specific relationships between branch lengths (for example, that the most recent branch is longer than all the others). The performance of the method is evaluated against simulated data and is also applied to historic mitochondrial data from the Nuu-Chah-Nulth people of North America. The method can be used to obtain estimates of the TMRCA of the sample. The relationship of these estimates to those given by “Thomson’s estimator” is explored.

## INTRODUCTION

The pattern of genetic variation in contemporary populations is the product of several factors including the population demographic history, mutation and adaptation to an organism’s environment. Genetic variation data has been successfully used in areas as diverse as tracing human prehistory and in identifying the origins both of genetic diseases and conditions that may confer immunity to them. New advances continue to be made powered by increases in both the availability of genomic data and in computational power. However, the view we obtain of the past in this way is necessarily obscured by the stochastic nature of the processes that created the data. Further, the validity of our inferences is dependent on the assumptions underlying the models that are used to link population history to observable data.

In recent times, the central component of most such models has been the coalescent tree associated with a genomic sample. Such a tree represents a genealogical history with attributes such as branch lengths, coalescence times and the time to the most recent common ancestor (TMRCA) as well as topological and combinatorial properties (for a summary see Simon and Huttley, 2021). Here we describe a method for estimating branch lengths or, equivalently, coalescence times from an observed site frequency spectrum (SFS) in the absence of selection and recombination. This method differs from previous approaches in not requiring an assumption that the data was produced by some particular known model or demographic history. Freedom from such assumptions is valuable, given that the relationship of the coalescence times itself provides information as to the demographic history of the population. For example, if a population has experienced expansion and the sequences considered have not been influenced by natural selection, the branches closest to the present will be relatively long compared to those close to the root, i.e. the tree will be more “star-shaped” (Hein et al., 2005).

An estimate for the coalescence time for the case of *n* = 2 sequences was given by Tajima (1983) (see also Tavaré et al., 1997), using analytic Bayesian methods. In this simple case, there is only one coalescence time, which is also the time to the most recent common ancestor (TMRCA), and the SFS is equivalent to the number of segregating sites. From a Bayesian perspective, Tajima uses the Wright-Fisher model to derive the prior distribution, although both Tajima and Tavaré et al. appear to have regarded the problem in terms of estimation of the coalescence times for a specific sample resulting from evolution under a Wright-Fisher model. Such an analytic method could not be extended directly to larger sample sizes, because this would involve multiple possible forms of the genealogical tree. Further advances on the problem made use of more computation-intensive Monte Carlo methods. These involved repeated simulations of evolutionary models, allowing various population genetic statistics to be calculated for each sample tree generated by the simulation and then averaged over the set. Tavaré et al. (1997) combined simulation with a rejection algorithm to provide a method of estimating individual branch lengths from an observed number of segregating sites on the assumption that the population was generated by a Wright-Fisher model. This method had the benefit of relative simplicity, but did not use all of the information contained in the SFS. A more complex approach based on importance sampling was used in Griffiths and Tavaré (1994). This made use of the variant data at the level of the individual sequences in the observed alignment, which provides more information than the summary given by the SFS. However, this method also depended on the assumption that the population had constant size or a specific known demographic history. It involved sampling genealogies consistent with the data by a backward in time process of generating successive mutation and coalescence events. Quantities such as branch lengths were then averaged over these sample genealogies (see also Felsenstein et al., 1999; Stephens and Donnelly, 2000; Wakeley, 2009). Other researchers used Markov Chain Monte Carlo (MCMC) methods to estimate other scalar parameters such as effective population size and mutation rate (Kuhner et al., 1995) and the time since a mutation event associated with a polymorphism (Markovtsova et al., 2000). These methods also assumed that the data was produced by a Wright-Fisher model.

A significant technical advance was made by the introduction of simulation software packages that could implement more complex evolutionary models, the most widely used being Hudson’s `ms` (Hudson, 2002). These allowed rejection algorithms and related approximate Bayesian computation (ABC) methods to be used to estimate parameters associated with more complex models incorporating selection and recombination. Examples include estimation of the time since the fixation of an allele under positive selection (Przeworski, 2003) and the time of origin and associated selection coefficient of such an allele (Peter et al., 2012). These methods are more suitable for problems involving smaller sets of scalar parameters and a relatively small number of scalar statistics summarising the observed data. However, simulation software itself is now indispensable in assessing the effectiveness of any inferential method in population genetics.

Some previous methods also addressed the question of estimating coalescence times without making assumptions as to the underlying evolutionary model and are therefore often referred to as “model-free” methods. Thomson et al. (2000) proposed as an estimator of the TMRCA of a sample the average number of mutations segregating in the sampled sequences divided by the mutation rate. This estimator is unbiased and requires no assumptions as to the demographic history of the sample. A closely related “model-free” estimator for the TMRCA, which does not, however, share the property of being unbiased, was proposed by Tang et al. (2002). It requires knowledge of the sizes of the two clades formed by the basal split of the tree. A “model-free” approach that estimates all coalescence times of a sample, rather than the TMRCA alone, was proposed by Meligkotsidou and Fearnhead (2005). This method requires that the full genealogy (tree topology) of the sample and the number of mutations on each branch are known. To address the question of how the specific tree topology is to be obtained, the authors cite the method of Gusfield (1991). However, this solution only obtains an instance of a tree that could have generated the data, which may not be the true tree or even a likely estimate of it. A likelihood-based method of estimating tree topology would generally involve calculating the joint maximum likelihood of the topology and the branch lengths, as is done in phylogenetic inference. This would make a further method to estimate branch lengths redundant. On the other hand, Thomson’s estimator does not require any knowledge of tree shape. Unlike our method, Thomson’s estimator does not extend to the estimation of all *n −* 1 coalescence times associated with a sample of size *n*. However, since the TMRCA is the final coalescence time, it provides a critical comparison to our method.

A key development subsequent to most of the papers cited above is the availability of powerful open source general-purpose MCMC software packages for Bayesian inference, which facilitate modelling with a wide range of discrete, continuous and multivariate probability distributions. We have made use of one of these software packages to demonstrate a general method for estimating the joint posterior distributions of the branch lengths of a genealogical tree, or the equivalent coalescence times, from a sample SFS, under the assumption of selective neutrality. The key advantage of this approach over previous methods is that it does not require that the evolutionary process that generated the tree, or the tree topology, be known. To achieve this, a novel uninformative prior distribution on branch lengths / coalescent times is proposed (although informative priors can also be designed). The fact that a Bayesian joint posterior distribution can be sampled has benefits in its own right in that it allows probabilities to be calculated for other hypotheses: for example, that one particular branch length is greater than another or that the sample had a given number of extant ancestors at some point in the past. The degree of uncertainty associated with estimates due to the stochastic nature of mutation and uncertainty as to tree shape can be visualised in terms of the “spread” of the posterior distributions. The method is also capable of being extended to accommodate the possibility of errors having been introduced into the data by upstream processes such as sequence read mapping and sequencing. Lastly, the method has the benefit of simplicity. This is primarily due to the use of the matrix representation of the combinatorial structure of genealogical trees, which simplifies both the sampling of trees and the computation of likelihood.

We test the method using coalescent simulations arising from different demographic scenarios. This demonstrates that valuable information about coalescence times can be derived from SFS data in the absence of assumptions about demographic history. We also show that significant improvements in inference can be achieved by using an appropriate informative prior. The degree of improvement is influenced by the demographic history of the population. The uncertainty in these inferences due to stochastic factors can be visualised and quantified.

## MATERIALS AND METHODS

The methods used in this paper draw from the theory set out in Simon and Huttley (2021). The focus is again be on the genealogy of a genomic sample conceptualised as a coalescent tree, and on the SFS as a summary of the observed sample data. The genomic tree for a sample of *n* sequences will again have a set of branch lengths *t*_2_, … *t*_*n*_, measured in generations. The total tree length *T* is the total time spent in the tree by all sequences since the most recent common ancestor of the sample:

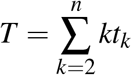

Our aim is to generate samples from the posterior joint distribution of branch lengths relating to the genealogy. We address this problem in two parts: estimation of total tree length *T* and estimation of the relative branch lengths (*t*_2_*/T*,… *t*_*n*_*/T*). We develop a Bayesian inferential model in each case, sample from the posterior distributions of each model using MCMC and combine the results to obtain a sample from the posterior distribution of branch lengths.

### Estimating total tree length from the number of segregating sites

We will begin by defining a simple Bayesian model for the estimation of *T*. This can be considered as one way of generalising the problem originally considered in Tajima (1983). The SFS for a sample of *n* aligned sequences is defined as the vector **s** of numbers *s*_*i*_ for *i* = 1, …, *n −*1, where *s*_*i*_ is the number of segregating sites at which the mutant allele occurs in *i* members of the sample. The total number of segregating sites *S*_*n*_ is then given by:

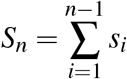

It is again assumed that mutations occur as a Poisson process, that is, the number of mutations occurring in a single branch is given by a Poisson distribution with parameter given by the product of the branch length and the per generation sequence mutation rate *µ*. Let us initially assume that *µ* is known. Then the probability of an observed value of *y* for *S*_*n*_ is given by the Poisson distribution with parameter *µT* :

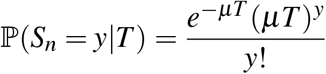

We will use a gamma distribution Gamma(*k, θ*) as a prior distribution for *µT* as it is the conjugate prior to the Poisson distribution (Gelman et al., 2014, §2.6) (we use the shape / scale convention for the gamma distribution). For an uninformative or diffuse prior distribution we will use Gamma(1, ∞). For this prior, the mode of the posterior distribution of the Poisson parameter is equal to the maximum likelihood estimate. Applying the method of conjugate priors (Gelman et al., 2014) then gives the following result for the posterior distribution of the parameter *µT* :

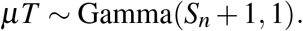

Using the scaling property of the gamma distribution gives the following posterior distribution for *T* :

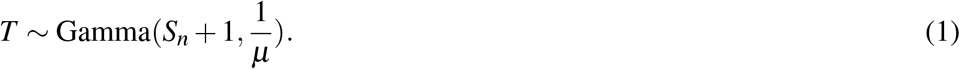

This posterior distribution has been described analytically, but we will use an MCMC model in order to incorporate uncertainty in mutation rate. There are two sources of uncertainty inherent in the use of an estimate of mutation rate. One is the sampling error associated with the estimation of the average mutation rate over the entire genome. Often, a more significant issue is the variance in mutation rate along the genome of humans and other organisms (Hodgkinson et al., 2009; Simon and Huttley, 2020), that is, the variance in mutation rate between segments of a given length. Our model using an uninformative prior on total tree length will incorporate uncertainty about the mutation rate by modelling it by a beta distribution with mean and standard error supplied to the model. A beta distribution is chosen as it takes values on the unit interval and has two parameters (c.f. Dunson and Tindall, 2000). It is also the conjugate prior of the binomial distribution, which is the natural form of the likelihood of a binary probability (or frequency) conditioned on sample data. Later in this Section, we also describe the use of an informative prior distribution for total tree length. This allows a uniform prior distribution on the mutation rate to be used.

### Inferring relative and absolute branch lengths

We again consider a sample of *n* sequences subtending a genealogical tree and this time seek to infer the joint posterior distribution of the relative branch lengths from the SFS of the sample. In equation (8) of Simon and Huttley (2021), the likelihood function for the SFS was given by:

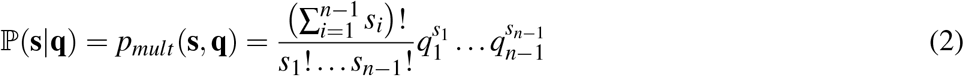

where **q** is a vector of probabilities given by:

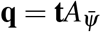

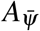 is the matrix representation of the tree partition class 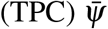associated with the genealogical tree *Ψ* subtending the sample; **t** is the vector of relative branch lengths given by:

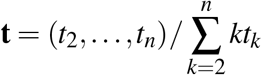

and *p*_*mult*_ is the probability mass function of the multinomial distribution.

It is also necessary to define a prior distribution for the estimation of relative branch lengths. The prior distribution will be defined over the joint parameters of TPCs and vectors of relative branch lengths, the latter denoted by **t**. As in Simon and Huttley (2021), the parameter space for TPCs is 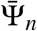 with the Equal Rates Markov (ERM) measure and the parameter space for relative branch lengths is the (*n−* 2)-dimensional simplex in 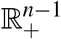 given by

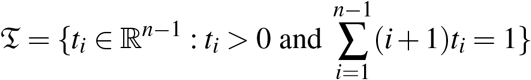

The model parameter space for the inference of relative branch lengths is the Cartesian product space 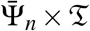. The prior distribution on this space will be the Cartesian product of two independent distributions defined on 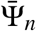 and 𝔗 respectively. The assumption that these distributions are independent reflects our lack of knowledge of any correlation between TPCs and relative branch lengths. The prior distribution on 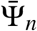 will be a categorical distribution with parameters given by the probabilities of each TPC under the ERM measure.

The uninformative prior distribution on 𝔗 will be the uniform distribution on this space, which we denote *𝒲*. In Simon and Huttley (2021) the linear bijection *J*_*n*_ from Δ^*n−*2^ to 𝔗 was defined as the diagonal matrix whose diagonal elements are 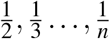. Since the linear transformation of a uniform distribution is a uniform distribution (Rudin, 1987), one can sample from *𝒲* by sampling from the uniform distribution on Δ ^*n−*2^ and transforming by *J*_*n*_. (The uniform distribution on Δ ^*n−*2^ is a Dirichlet distribution with all parameters set to 1.) The mean of this prior distribution on 𝔗 is therefore given by:

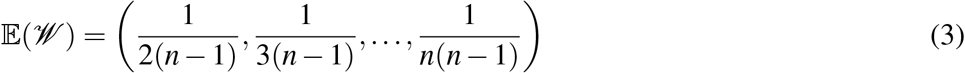

Having thus defined a likelihood and (joint) prior distribution for our model, MCMC software can be used to sample from the joint posterior distribution of relative tree lengths and TPCs, given the SFS data. The TPC parameter is “integrated out” by considering the branch length samples only, resulting in a marginal posterior distribution for branch lengths (for marginal posterior distributions see Gelman et al., 2014, Chapter 3). This posterior distribution can then be multiplied with the posterior distribution of total tree length. This results in what we can call a convolution posterior distribution of absolute branch lengths. We will however, simply use the term “posterior distribution”, unless we wish to draw attention to its derivation. Implementation uses the general-purpose MCMC software package PyMC3 (Salvatier et al., 2016) and the relevant Python code is provided in a github repository at https://github.com/helmutsimon/bayescoalescentest. Bayesian point estimates can be derived in the usual fashion as the expected value of the posterior distribution, known as the minimum mean square error (MMSE) estimate as it minimises the mean squared error relative to that posterior distribution (Jaynes, 2003, p. 172).

To sample tree matrices from 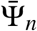 according to the ERM measure, we use the theory set out in Simon and Huttley (2021) in which tree matrices represent sequences of unordered partitions. A sequence of ordered partitions can be pictured as being generated by the division of a set of *n* objects into *n* singletons by successively replacing the spaces between the objects (initially there are *n −* 1 of these) by dividers. The elements of the sequence can be represented by integer indices indicating which of the remaining gaps is filled at each step. In this representation, known as the Lehmer code (Lehmer, 1960; Devroye, 1986, Chapter 13 p. 644), the first index is in the range 1, …, *n −* 1, the second in the range 1, …, *n −* 2 and so on. Each index in the sequence represents the branching event at level *k*, starting at *k* = 2, conditioned on the previous numbers in the sequence. We can therefore sample uniformly from sequences of ordered partitions by a sequence of independent categorical distributions with *n −* 1, *n −* 2, …, 2 categories. These sequences of ordered partitions can be transformed into TPCs by forgetting the ordering of the partitions, thus sampling from 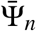 according to the ERM measure. This allows the use of a Metropolis-within-Gibbs MCMC proposal/acceptance method (implemented in PyMC3 as `CategoricalGibbsMetropolis`), which samples from each of the categorical distributions independently. Use of this proposal/acceptance method achieves convergence of the Markov chain for larger sample sizes. The posterior distributions of these indices can be interpreted in terms of the branching events in the tree. This is clearest in the *k* = 2 case of *n −* 1 categories, which represents the basal split into two clades from the root of the tree.

### Inference of derived quantities

The MCMC process defined above provides us with samples from the joint posterior distribution of branch lengths. These can be used to obtain posterior distributions for other related quantities. The joint posterior distribution of coalescence times (viewed back in time from the present) can be obtained as cumulative sums of branch lengths. Note that there is a bijective linear transformation that takes the ratios of branch lengths to total tree length to the ratios of coalescent times to total tree length, which could have been used to create a slightly more complex model to estimate relative coalescence times directly.

The last or (*n −* 1)^*st*^ coalescence time is the sum of all branch lengths and hence equal to the TMRCA. By restricting our attention to this last coalescence time, we obtain a marginal posterior distribution for the TMRCA. We can use the expected value of this marginal distribution as a point estimate of the TMRCA, which is an MMSE estimator relative to the marginal distribution.

We can also use the posterior distribution of branch lengths to calculate what we refer to as the ancestral probabilities associated with a specific genomic sample *X*. This is the probability ℙ (*A*_*X*_ (*t*) = *k*) for 1 *≤k ≤n*, where *A*_*X*_ (*t*) is the number of ancestors of *X* at time *t*. ℙ (*A*_*X*_ (*t*) = *k*) can be estimated by counting the number of variates in the sample from the posterior distribution of coalescence times for which *t* lies between the relevant coalescence times. ℙ (*A*_*X*_ (*t*) = *k*) can be considered as a Bayesian analogue of the “ancestral process” of classical population genetics (Tavaré, 1984; Griffiths and Tavaré, 1998), denoted by ℙ (*A*_*n*_(*t*) = *k*) where in this context *A*_*n*_(*t*) denotes the number of ancestors of a sample of size *n* at time *t* and the probability relates to a specific evolutionary model, which is usually the Wright-Fisher model or a modification involving some deterministic variation in population size. Here ℙ (*A*_*n*_(*t*) = *k*) is the probability that *A*_*n*_(*t*) = *k* for a random run of the evolutionary process. It might be determined experimentally by multiple simulations of the process. In a similar way, one can determine the probability (in Bayesian terms) of other hypotheses. For example, one could count the MCMC variates to determine the probability that one branch length *t*_*i*_ is greater than another (*t* _*j*_), or that a particular branch is the longest.

### Model-based prior distributions

In the previous sections, uninformative uniform prior distributions were employed in the Bayesian models used to estimate relative and absolute branch lengths. This is a reasonable choice if one has no prior knowledge as to the evolutionary process that produced the data. However, we may have a prior belief that the population evolved according to a particular demographic model, such as constant or exponentially increasing population size. Such a model will define a probability distribution on the space 𝔗 of relative branch lengths, which we will denote by *G*. A prior belief in this evolutionary model would be reflected by using *G*, rather than the uniform distribution on 𝔗, as a prior distribution for the analysis of relative branch lengths in the MCMC model. Although *G* cannot typically be expressed in terms of standard probability distributions, one can use an approximation to *G* as a prior distribution, by matching the expected value vector and covariance matrix of *G*. To do this, a coalescent simulation package is used to generate samples from *G*. These variates are firstly transformed into the standard simplex Δ_*n−*2_ using the linear transformation 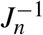 and then into variates in ℝ^*n−*1^ by means of a log-ratio transformation as used in the statistical analysis of compositional data (Aitchison, 1982) and implemented as the `StickBreaking` transform in PyMC3. The transformation used in calculations for this paper is:

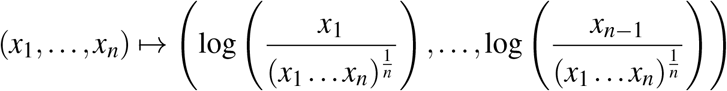

The sample expectation vector and covariance matrix of these transformed variates in Euclidean space are then calculated and used to define a multivariate normal (MVN) distribution with these parameters. This MVN distribution is used as the basis of the prior distribution by reverse transforming the variates back into 𝔗.

The expected mean and standard deviation of the total tree length can also be derived from simulations of the model. These can be used as parameters defining the gamma prior distribution for total tree length.

### Data

Empirical data pertaining to the Nuu-Chah-Nulth people was obtained from Griffiths and Tavaré (1994, Table 1).

**Table 1.** Comparison of the use of uniform and MVN prior distributions to estimate branch lengths and TMRCA using Bayesian inference for a range of models. The first two columns show the sample size (*n*) and mean number of segregating sites 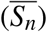. The other columns show the MSE for branch lengths and TMRCA under the two priors and for the TMRCA under Thomson’s estimator. (a) A constant size population model. Errors are shown in units of 10^9^ generations squared. (b) A constant size population model with a higher mutation rate. Errors are shown in units of 10^9^ generations squared. (c) An exponentially increasing demographic model. Errors are shown in units of 10^4^ generations squared. In each instance 100 trials were used.

| n | $\bar{S}_n$ | Branch lengths | | TMRCA | | |
| --- | --- | --- | --- | --- | --- | --- |
|  |  | Uniform | MVN | Uniform | MVN | Thomson |
| 10 | 10.1 | 17.5 | 13.5 | 11.4 | 10.9 | 13.0 |
| 25 | 14.3 | 32.0 | 16.3 | 32.7 | 16.3 | 23.6 |
| 50 | 16.2 | 66.1 | 23.0 | 74.2 | 19.7 | 22.5 |
| (a) Growth rate = 0, $\theta = 3.6$ | | | | | | |
| 10 | 30.86 | 10.2 | 8.45 | 6.04 | 5.20 | 6.86 |
| 25 | 40.51 | 11.2 | 10.4 | 9.41 | 5.58 | 5.85 |
| 50 | 49.51 | 39.3 | 13.6 | 38.3 | 7.99 | 8.22 |
| (b) Growth rate = 0, $\theta = 10.8$ | | | | | | |
| 10 | 11.4 | 14.8 | 3.74 | 20.3 | 0.40 | 6.40 |
| 25 | 21.6 | 10.2 | 2.72 | 39.6 | 0.57 | 10.1 |
| 50 | 45.4 | 5.33 | 1.90 | 36.4 | 0.55 | 7.78 |
| (c) Growth rate = $10^{-2}$, $\theta = 800$ | | | | | | |

### Software

Scripts and Jupyter notebooks developed for this work were written in Python version *≥*3.5 and are freely available under the General Public License at https://github.com/helmutsimon/bayescoalescentest.

Synthetic data was generated using *msprime* (Kelleher et al., 2016), which is a Python library implementing a version of the Hudson `ms` (Hudson, 2002) coalescent simulation software. Each *msprime* simulation yields a genealogical tree with mutations added. A range of parameters can be set, in particular, a wide range of demographic histories. The resultant site frequency spectrum, the coalescence times and the tree partition class can be determined from the tree objects returned. Inferred branch lengths or other parameters can thus be compared to the actual values resulting from a specific simulation.

Dependencies included `cogent3` 2019.12.6a (Knight et al., 2007), `pyMC3` 3.11.0 (Salvatier et al., 2016), `theano` 1.0.4 (Al-Rfou et al., 2016), `arviz` 0.11.1 (Kumar et al., 2019), `scipy` 1.2.1 (Virtanen et al., 2020), *msprime* 0.7.0 (Kelleher et al., 2016), `numpy` 1.16.3 (Virtanen et al., 2020), `pandas` 0.24.2 (McKinney, 2010), `more-itertools` 8.7.0 (Rose and Bayles, 2012), `click` 6.7 (Ronacher, 2009), `scitrack` 0.1.3 (Huttley, 2016), `matplotlib` 3.0.3 (Hunter, 2007) and `seaborn` 0.9.0 (Waskom et al., 2017).

### Data availability statement

The authors state that all data necessary for confirming the conclusions presented in the article are represented fully within the article. Supplementary Information, data sets, run logs and scripts produced for this work are available at Zenodo https://zenodo.org/record/5121182, doi 10.5281/zenodo.5121182 under the Creative Commons Attribution-Share Alike license.

## RESULTS

### Testing inference using simulations with a uniform prior

we first illustrate the method by using synthetic data generated by *msprime* for a range of different sample sizes and demographic scenarios in Figure 1. In these cases the sample size *n* = 5 was used. We can directly compare the posterior distributions for coalescence times to the actual coalescence times for a single simulation run. Figure 1a shows coalescent times from a simulation with constant population size, Figure 1b from a population experiencing exponential growth (at a rate of 10^*−*4^ per generation) and Figure 1c from a more complex demographic scenario of constant population size (going forward in time) followed by a population bottleneck followed by exponential growth (see Supplementary Information Section 1 for details). In each figure, the posterior distributions of coalescence times are plotted for *k* = 1, …, *n−*1 and the actual coalescence times are shown by vertical lines. The “spread” of the distribution reflects uncertainty due to the stochastic nature of mutation and the unknown tree structure. In Figure 1b, one can observe significant underestimation of the first two coalescence times. We will discuss possible reasons for this below.

**Figure 1.**
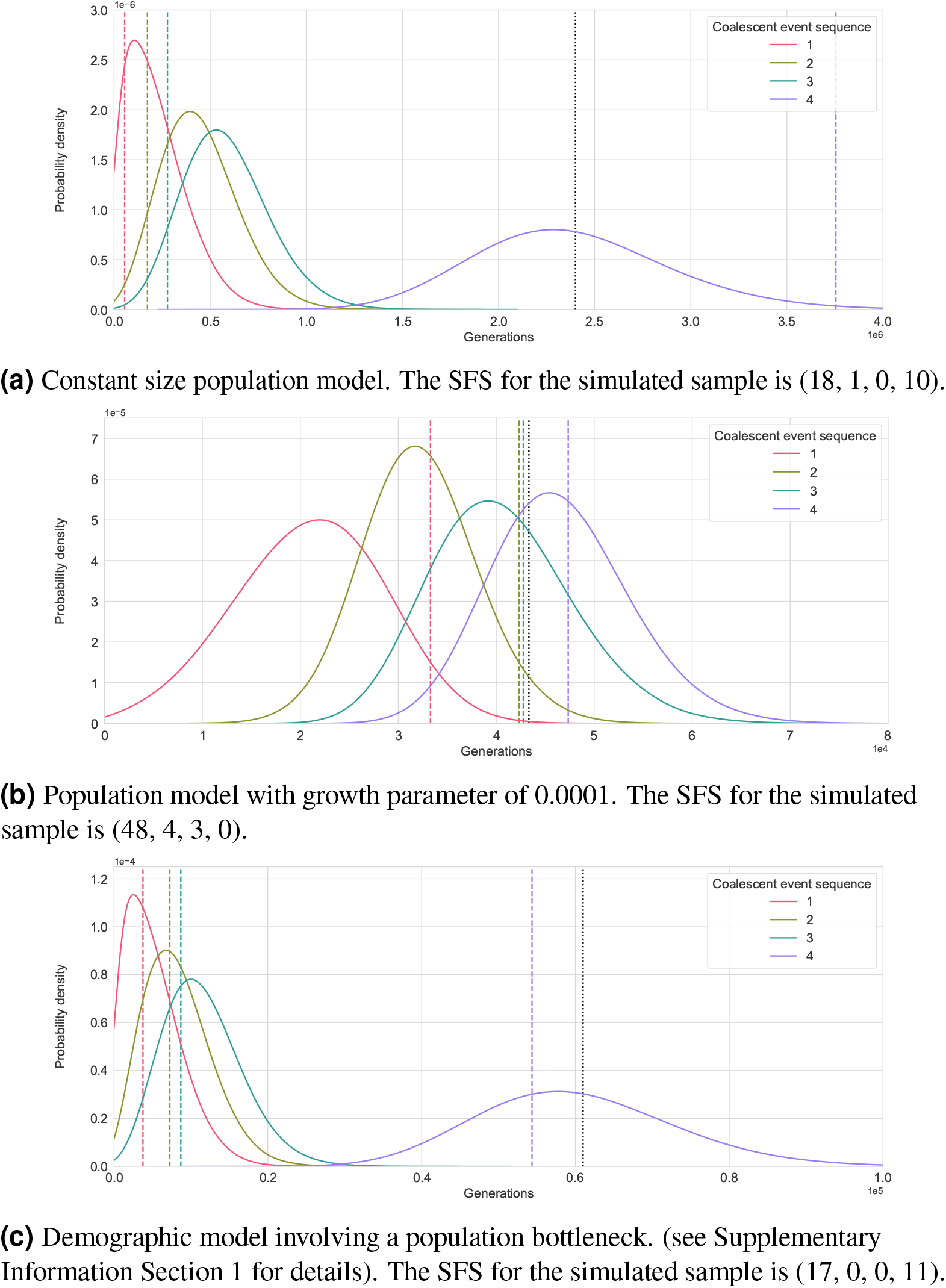
Posterior density functions of coalescence times obtained from simulated samples of size *n* = 5 using different population models. The actual coalescence times being estimated are shown as vertical lines. The numbers in the legend represent the order of coalescence times counting backward from the most recent coalescent event to the most distant (the TMRCA). The Thomson estimate for the TMRCA is shown as a black dotted line.

These examples also illustrate the estimation of the TMRCA. The “highest” coalescence time (labelled *k* = *n −* 1) is also the TMRCA, and Figures, 1a and 1c show quite good agreement between the marginal posterior distribution of the TMRCA and the true values. The posterior distribution also provides an indication of the accuracy that can be expected from these estimates. The TMRCA itself can be estimated with less computational effort using Thomson’s estimator (Thomson et al., 2000), which is shown as a dotted black line on these figures. In Figure 1a the MMSE estimate of the TMRCA is not in good agreement with the true value, although it is close that obtained using Thomson’s estimator.

A set of figures using the same population models, but a sample size of *n* = 8 are shown in Supplementary Information Section 2. These show improved estimates for the constant size and exponentially increasing population models in particular, possibly due to a higher *S*_*n*_.

### Simulations using model-based priors

One object of using an uninformative prior distribution in Bayesian inference is to maximise the influence of the observed data on the outcome relative to that of the prior distribution. While the uniform Dirichlet prior distribution defined on relative branch lengths is intended to be uninformative, it does have an expected value whose components have the relative proportions 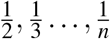 (equation 3). That is, the mean branch lengths decrease as *k* increases from 2 to *n*, albeit more slowly than for the Wright-Fisher distribution. This characteristic of the prior distribution has the potential to bias results. An example was provided by Figure 1b, which analyses a sample of size *n* = 5 from a simulated population undergoing exponential growth. In such a case, the coalescence times nearest to the present are expected to be relatively large, contrary to the case for the uniform prior distribution. Figure 1b shows that the first two coalescence times were significantly underestimated.

We investigated this matter further by comparing the use of a uniform (uninformative) prior to a MVN (model-based) prior also taking account of sample size and the number of segregating sites. We selected a number of models, with various sample sizes, mutation rates and population growth rates, and ran 100 simulations for each. The resulting samples of relative branch lengths were first used to derive the parameters for the MVN prior using the method in the sub-section *Model-based prior distributions* above. The MCMC method was then used to derive point (MMSE) estimates of the vectors of branch lengths from the SFS produced by each of the 100 simulations, using both choices of prior distribution. The accuracy of each estimate relative to the true values was measured by the mean squared error (MSE), taken over the 100 simulations. That is, a quadratic loss function was selected for the comparison. This choice was made to be consistent with general practice and the use of MMSE point estimates. The MSE was also calculated for estimates of the TMRCA, including estimates made by Thomson’s method. The results are shown in Table 1. Subtable 1a shows results for a population with a constant population size and a fixed mutation rate. We see that for each of the sample sizes shown the MSE with the MVN prior is lower than that with the uniform prior, but the margin is more pronounced for the larger sample sizes. Subtable 1b shows results for the same constant population size evolutionary model as for Subtable 1a, but with 3 *×* mutation rate. The errors are lower than in Subtable 1a for all sample sizes and both prior distributions. Subtable 1c shows results for a different evolutionary model, namely a population with a relatively rapid growth rate of 10^*−*2^. We see that the reduction in MSE using the MVN prior compared to the uniform prior is more dramatic than for the constant size population.

The Thomson estimates of the TMRCA are better than those using the uniform prior, possibly excepting the smallest sample sizes. The relative performance of the MVN prior and the Thomson estimator vary with mutation rate and evolutionary model. In the case of an expanding population, the MVN prior strongly outperforms Thomson.

To further illustrate the effect of using an MVN prior for an expanding population, the analysis illustrated in Figure 1b was rerun using a prior distribution based on the correct model population growth rate of 0.0001. The outcome shown in Figure 2 is in substantially better agreement with the true values than was obtained using the uniform prior as in Figure 1b.

**Figure 2.**
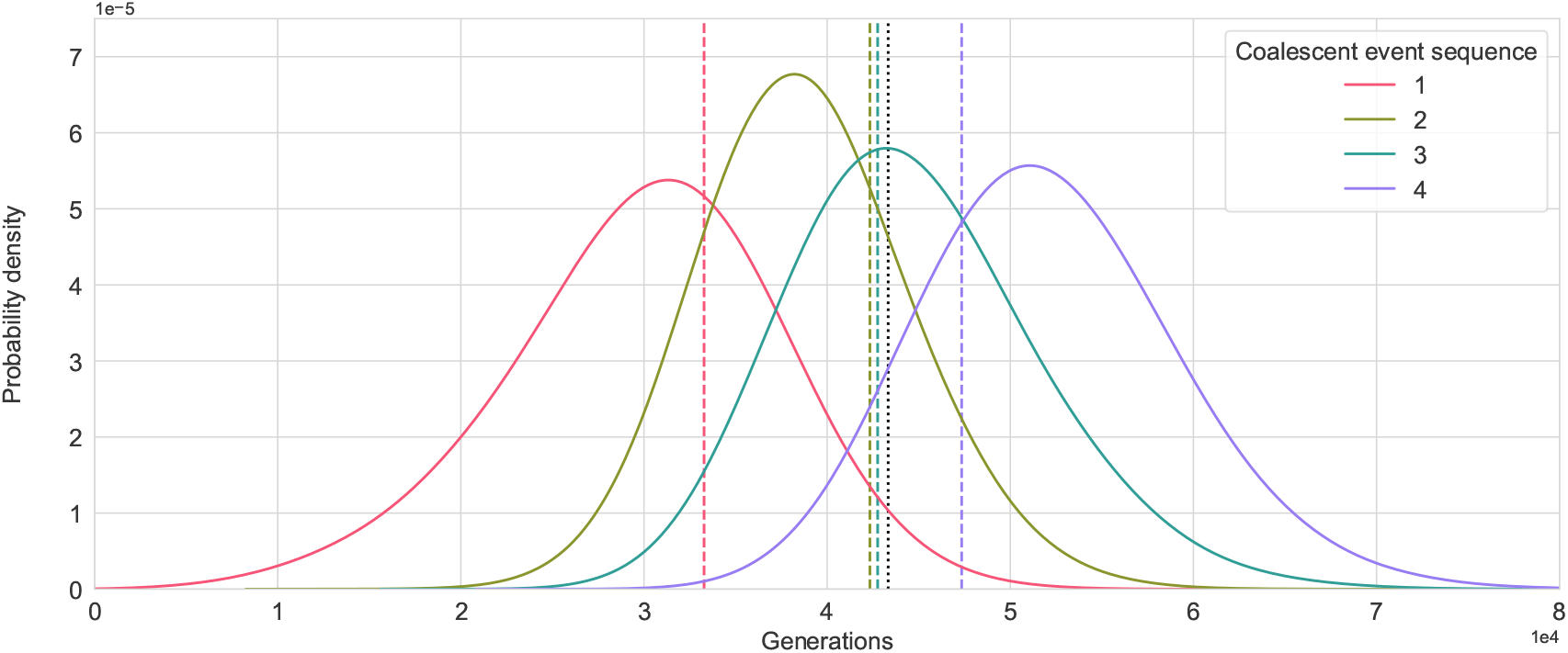
Derived posterior distributions for coalescence times using the same data as in Figure 1b and a posterior distribution based on a population growth rate of 0.0001, consistent with the model that produced the data. Actual values are shown as vertical lines. The numbers in the legend represent the order of coalescence times counting backward from the present.

### Data from the Nuu-Chah-Nulth population

We analysed mitochondrial DNA from the native American Nuu-Chah-Nulth people of Vancouver Island using a classic data set that originally appeared in Ward et al. (1991) and has been the subject of several studies (Griffiths and Tavaré, 1994; Kuhner et al., 1995; Marjoram and Tavaré, 2006). This was undertaken to demonstrate the application of the method to empirical data and to determine the extent to which our results agreed with previous analyses using different methods.

Griffiths and Tavaré (1994) considered 55 samples of a mitochondrial sequence of 352 bps, containing 18 segregating sites. They assumed, on the basis of archaeological evidence, that the female population had remained at a constant size of 600 for *>* 6000 years. The value of Tajima’s D for the data is -0.5, which is consistent with the assumption of constant population size. Basic results on the Wright-Fisher model (Hein et al., 2005) tell us that, without conditioning on the data, the expected TMRCA for the sample is 1178 generations and the expected total tree length is 5491 generations.Griffiths and Tavaré first estimate the sequence mutation rate and standard deviation, conditioned on the data and these demographic assumptions, to be 4.0*±*1.2 *×*10^*−*3^. We used a simpler model to estimate the mutation rate by MCMC, using only the total number number of segregating sites to represent the observed data. We used a prior distribution on total tree length as given by the Wright-Fisher model and a uniform prior distribution on mutation rate. This resulted in an estimate of 3.95 *±* 1.4 *×*10^*−*3^ for mutation rate, which is in good agreement with Griffiths and Tavaré. Griffiths and Tavaré then estimated the TMRCA of the sample conditioned on the data at 720 generations, using an importance sampling method (see also the discussion in Felsenstein et al., 1999). The estimate of the TMRCA using Thomson’s method together with Griffiths and Tavaré’s estimate of the mutation rate is 595.5 generations.

We analysed the data using two different prior distributions on relative branch lengths: an uninformative (uniform) prior and a prior based on the Wright-Fisher model. In the latter case, we used the MVN approximation method described in Materials and Methods. Both models involve simultaneous estimation of the branch lengths and the mutation rate, so an informative prior is required for one of these quantities. When using a uniform prior on relative branch lengths, we used a beta prior distribution for mutation rate with parameters derived from our estimate of 3.95*±*1.4*×*10^*−*3^ for the mutation rate. When using an MVN prior on branch lengths, we used a uniform prior for the mutation rate.

The results obtained using a uniform prior distribution on relative branch lengths are shown as a heatmap in Figure 3a. The heatmap illustrates the probability distribution function for each coalescence time. The probability distribution function values are normalised relative to their maximum for visual clarity. Such a heatmap representation is more intelligible for large sample sizes than the line plots shown earlier. In this case, the MMSE estimate of the TMRCA is 533.2 generations, the MMSE estimate of the total tree length is 6583.4 generations and the MMSE estimate of the mutation rate is 3.37*±*1.3*×*10^*−*3^. The results obtained using an MVN prior are shown in Figure 3b. In this case, the MMSE estimate of the TMRCA is 766.8 generations, the MMSE estimate of the total tree length is 5095.3 generations and the MMSE estimate of the mutation rate is 4.04*±*1.5*×*10^*−*3^. As might be expected, the estimate of the TMRCA obtained using a prior based on the Wright-Fisher model is closer to that obtained by Griffiths and Tavaré using a method that assumed this model.

**Figure 3.**
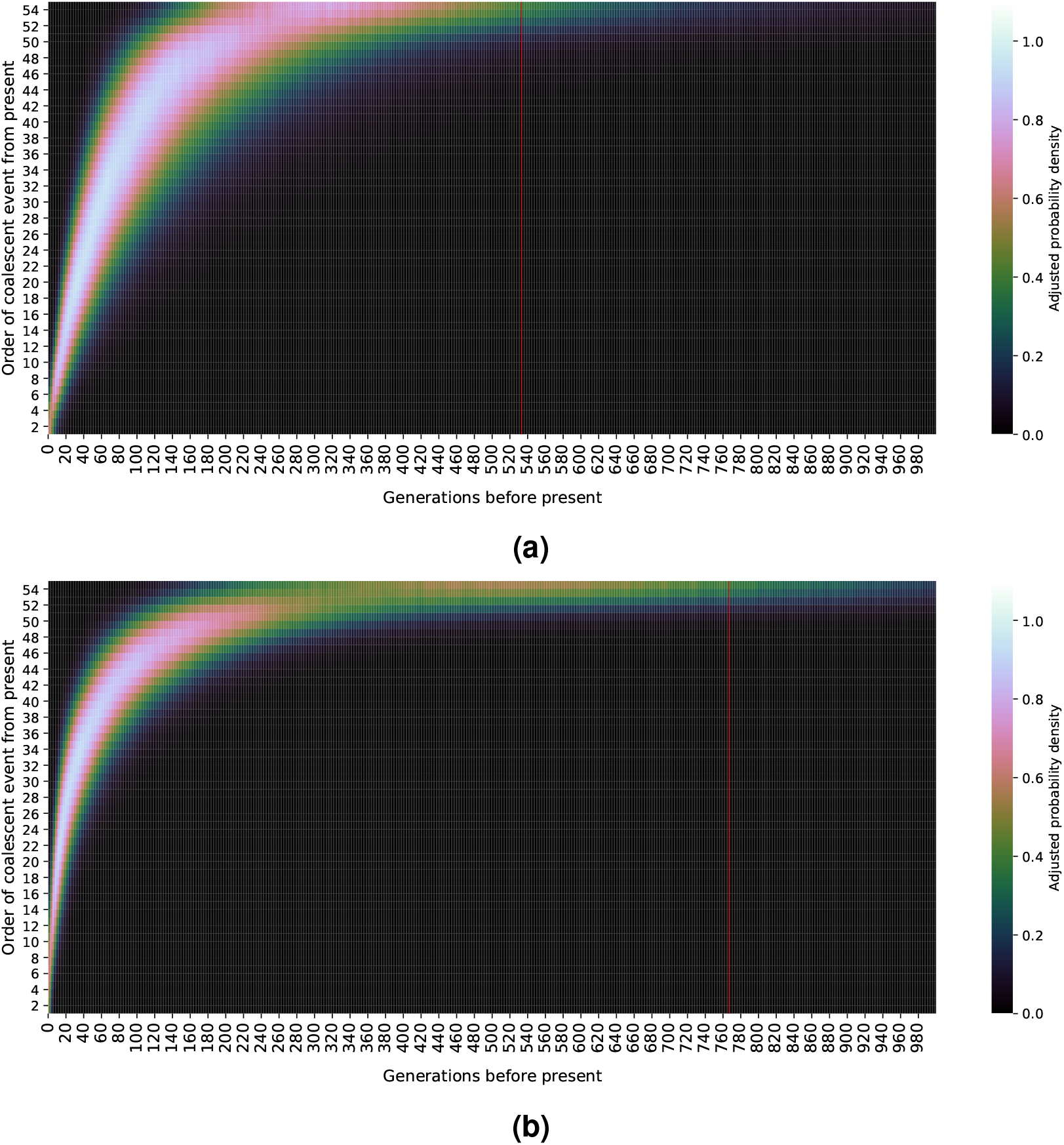
A heatmap representation of the posterior distribution of coalescence times estimated from 55 mitochondrial samples from the Nuu-Chah-Nulth people. The posterior mean estimate of the sample TMRCA is shown by a vertical red line. (a) An uninformative (uniform) prior distribution was used. (b) An multivariate normal prior distribution derived from the Wright-Fisher model was used.

In order to investigate the varying influence of the uniform prior distribution on different sample sizes, we also analysed a random sub-sample of size 12 from the full sample of 55 using this prior distribution. In this case the MMSE estimate of the TMRCA was 653.3 generations and the MMSE estimate of the mutation rate was again 3.37*±*1.3*×*10^*−*3^.

As stated in Materials and Methods, the MCMC model samples tree matrices by sampling from sequences of ordered partitions using a sequence of independent categorical distributions. These distributions represent branching events at each level of the tree. Therefore the MCMC results also give information about the tree structure, most clearly the sizes of the 2 clades formed by the basal split at the root of the tree. For this sample the most likely basal split is 24/31 with *p* = 0.56. The next most likely are 19/36 (*p* = 0.19) and 12/43 (*p* = 0.17). (The probabilities we give are for unordered partitions: the ordered partitions 24/31 and 31/24, for example, are equivalent and expected to be equiprobable). See Supplementary Figure S4.

## DISCUSSION

### Previous use of Bayesian methods

Related problems for very simple cases have previously been addressed using analytic Bayesian methods. These include calculating branch lengths when the sample size *n* = 2 (Tajima, 1983; Tavaré et al., 1997; Walsh, 2001) and the case where *S*_*n*_ = 0 (Dorit et al., 1995; Donnelly et al., 1996). The example closest to our work is Tajima (1983). This considers the case *n* = 2 and uses a prior distribution for *T* given by the Wright-Fisher model, that is, the exponential distribution with parameter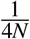. This is the same as the gamma distribution Gamma(1, 4*N*). Hence, using the method of conjugate priors, Tajima’s posterior distribution for the total tree length (twice the coalescent time) is, using our notation, given by:

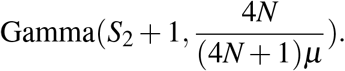

Comparing this result with equation (1) shows that in this case the use of different prior distributions makes little difference to the outcome, provided that the population size is reasonably large.

### Choice of prior distribution

Applications of Bayesian inference require explicit consideration of the choice of a prior distribution. The constant-size Wright-Fisher model may seem an obvious choice from which to derive a prior distribution. This choice is warranted if we have a reasonable prior belief that the size of the population has been constant over a period of time, as appeared to be the case in the Nuu-Chah-Nulth example. We have proposed the uniform distribution on the space of valid relative branch lengths as a preferable choice when no prior knowledge is available regarding the model that produced the data. If we have a prior belief that a given model may have produced the data, a prior distribution approximating the distribution of relative branch lengths associated with that model can be used. We have described a general method for approximating any model that can be simulated, by using a multivariate normal approximation transformed to the standard simplex. Other methods of approximation may also be effective, such as a kernel density estimate (KDE) using software such as the **KernelDensity** class in scikit-learn (Pedregosa et al., 2011). However, PyMC3 does not support this method for multivariate distributions.

We investigated the bias that can arise from use of the uniform prior distribution by comparing the outcomes with those of inferences for which the prior distribution is derived from an approximation to the simulation model. The results are shown in Table 1. Comparing each entry in Subtable 1b to the corresponding entry in Subtable 1a we see that, all other things being equal, a higher mutation rate, which leads to more segregating sites and hence more data, will result in better estimates. This is as expected. We further see that the relative advantage of an informative prior distribution relative to an uninformative prior generally decreases with more data (more segregating sites). This is also consistent with the principle that the relative influence of the prior will decrease with additional data. On the other hand, one can see a greater bias resulting from the uniform prior when the true distribution of branch lengths is very different to that under the uniform prior. This is illustrated for the case of a relatively high exponential growth rate of 10^*−*2^ (Subtable 1c), which results in the branch lengths for *k* = *n* being the largest on average, while mean branch lengths under the uniform distribution decrease as *k* increases.

Comparing results for different values of *n* in Subtable 1a, we see that the accuracy of estimates of branch lengths and of the TMRCA using the uniform prior decreases as *n* increases. We see the same effect, although to a lesser extent, in Subtable 1b. That the estimation of a one-dimensional population genetic parameter is subject to diminishing returns as sample size is increased is well known (Felsenstein, 2005). When estimating the joint probability distribution of branch lengths or coalescence times of genomic samples from a single population, the number of parameters being estimated also increases with sample size. Such an addition of new parameter dimensions through larger samples may negate any improvement in estimates unless the new samples add a sufficient quantum of new data (Gelman et al., 2014, §4.3). As the information available from a larger sample is spread over a larger number of parameter dimensions, the relative influence of the prior distribution may increase. The reason that increasing sample size does not have a direct effect on the amount of available data lies in turn with the correlation of the ancestries of the genomic sequences in the sample. For the Wright-Fisher model, the amount of additional information (the mean *S*_*n*_) only increases 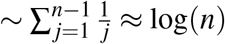 (Ewens, 2004, §9.6), which is not sufficient to compensate for the increase in the number of dimensions of the parameter set. The problem is inherited by marginal posterior distributions (Gelman et al., 2014, §4.3), such as the TMRCA in our case. A further example is provided by the Nuu-Chah-Nulth data set. In this case, the use of a sample size *n* = 55 resulted in a lower TMRCA estimate than for a sub-sample of *n* = 12, although a sub-sample cannot have a higher TMRCA than a full sample. However, this “curse of dimensionality” is not evident in the case of an exponentially increasing population in Subtable 1c. Here, increasing sample size appears to result in some degree of improvement in the accuracy of estimates, particularly in estimating joint branch lengths using the uniform prior. We can see from the second column of this Subtable that *S*_*n*_ increases approximately ∼ *n* and therefore the additional available data keeps pace with the dimensionality of the space in which it exists. This in turn results from the fact that in a population increasing in size, the ancestries of elements of the sample are not as highly correlated. The potential benefit or otherwise of increasing sample size can thus to some extent be predicted by the rate of increase in *S*_*n*_. It can also be seen from Table 1 that any negative effect of increasing sample size is less when a suitable prior distribution is used. This is to be expected, as the effect of the “curse of dimensionality” in these examples is to give greater weight to the prior distribution.

### Inference of derived quantities

A feature of this method is the inference of the joint posterior distribution of coalescence times or branch lengths. As stated previously, this can be used as the basis for computing other probabilities and distributions relevant to the sample. An example is given by the ancestral probabilities defined in Subsection *Inference of derived quantities* above. To illustrate, Figure 4, shows such ancestral probabilities derived from the posterior distributions for the data used in Figure 1b (exponentially increasing population size). Further illustrations of ancestral distributions are shown together with the relevant posterior distributions of coalescent times for simulations using sample size *n* = 8 in Supplementary Information Section 2.

**Figure 4.**
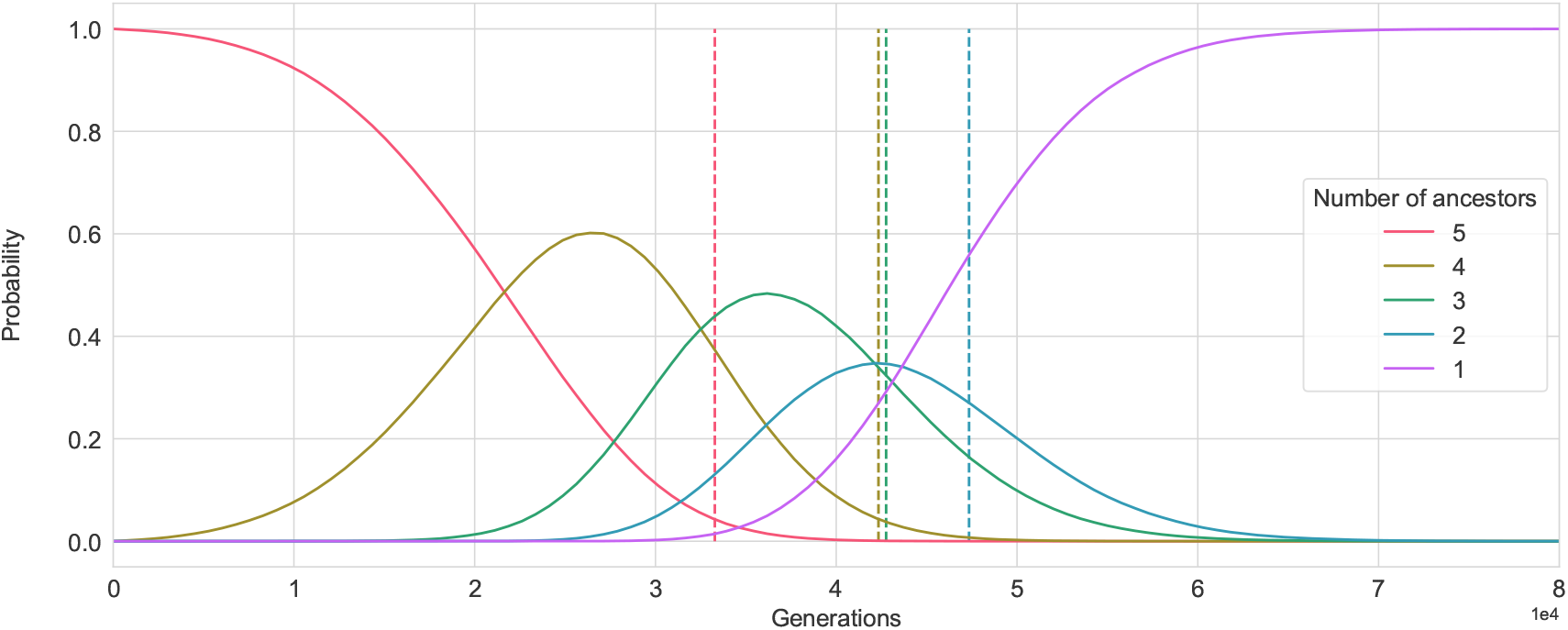
The ancestral probability distributions corresponding to the model and data used in Figure 1b (exponentially increasing population size). The plot for *k* = 1, …, 5 (Legend) shows the estimated probability that there are exactly *k* extant ancestors of the sample at the time *t* shown on the horizontal axis. The coalescent times, at which *k* ancestors of the sample reduce to *k −* 1 ancestors, are again shown as vertical lines.

### The Nuu-Chah-Nulth example

The practical effect of choice of prior distribution was illustrated in our analysis of Nuu-Chah-Nulth data, where both a uniform and (approximate) Wright-Fisher prior were used. The differing outcomes are exemplified by the respective point estimates for the TMRCA of 533 and 767 generations. The magnitude of this difference reflects the relative sparseness of the data (18 segregating sites occurring in 55 mitochondrial DNA samples). In fact, the estimate using the uniform prior was higher for a sub-sample of size 12 than for the full sample of size 55, as discussed above.

Some further observations can be made from our analysis. It can be seen from Figure 3 that the “spread” of the posterior distributions of coalescent times obtained was less when an informative prior was used, as one would expect. It can also be observed that Figure 3b shows characteristics more typical of a Wright-Fisher model, in that the coalescence times near the root of the tree are widely spaced, reflecting the relatively large inferred branch sizes for *k* = 2 and *k* = 3 in particular. We also saw that the most probable value for the basal split (24/31) had a posterior probability of 0.56. The fact that such a degree of uncertainty as to the basal split may exist needs to be considered when making population genetic inferences that rely on this knowledge (see, for example, Tang et al., 2002, referred to in the Introduction, and Yang et al. 2018).

We note that the Nuu-Chah-Nulth data is problematic due to the low population size, particularly relative to the sample size of 55. This means that the assumption common to the coalescent models used in this paper and by Griffiths and Tavaré that sample size sufficiently small relative to population size that more than one coalescence is unlikely in a single generation, does not apply.

### Thomson’s estimator for the TMRCA

We previously referred to Thomson’s estimator for the TMRCA (Thomson et al., 2000) as a precursor to “model-free” inference of population genetic parameters related to a sample of sequences. For a sample of *n* sequences, an SFS *s*_1_, …, *s*_*n−*1_ and a sequence mutation rate of *µ*, Thomson’s estimator is calculated by the following formula:

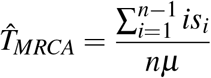

This expression has an intuitive interpretation as the average number of mutations in a sampled sequence, divided by the mutation rate *µ*, where only mutations that do not appear in the entire sample, i.e. those that are known to have occurred subsequent to the most recent common ancestor of the sample, are considered. The Thomson estimator is easily calculated and it is an unbiased estimator (Hudson, 2007). That is, it is unbiased relative to the probability distribution of mutations in a single genealogical tree. Calculation of the estimate does not depend on prior knowledge of properties of the genealogical tree such as the basal split (cf. Tang et al., 2002) or the tree topology (cf. Meligkotsidou and Fearnhead, 2005). However, properties of the estimator such as variance do depend on properties of the genealogical tree (Hudson, 2007). The application of Thomson’s estimator does require a prior estimate of the mutation rate. However, the method could no doubt be extended to quantify its sensitivity to the uncertainty which necessarily exists in any estimate of the mutation rate.

Our results have shown that Thomson’s estimator often gives a result similar to the mean of the posterior distribution for the TMRCA. The methods are in fact related in that they can both be derived from the same model. However, Thomson’s estimator is derived by the method of moments rather than Bayesian inference, as we show in Supplementary Information Section 4. It can be seen that Thomson’s method estimates the TMRCA directly, rather than as a sum of estimates of branch lengths, so the adverse effect of the “curse of dimensionality” is not as great as for the marginal Bayesian posterior distribution method. However, it is not clear from Table 1 whether larger sample sizes actually lead to significant improvement in Thomson estimates of the TMRCA and there may be scope for further theoretical work on this. Further, Subtable 1c shows that for some demographic histories, a strong choice of prior distribution can outweigh this benefit. Overall, it is clear that whenever the Bayesian method is applied, Thomson’s estimate should also be calculated for comparison.

### Why use the site frequency spectrum?

The SFS has been widely used as a summary statistic in population genetic inference (Nielsen et al., 2012; Han et al., 2013). In principle, more complete variant data could be obtained that identified, for every site segregating in the sample, the specific sequences in which the mutant allele appears. The likelihood function for such a set of full variant data depends not only on the TPC, but requires knowledge of the tree topology, as well as of relative branch lengths. Even for low sample sizes, where each TPC corresponds in a one-to-one fashion to a tree topology, the likelihood functions for the SFS and the full variant data generally differ and hence the posterior distributions of branch lengths will differ also. In other words, the SFS is not a sufficient statistic for inference of branch lengths. However, from a practical standpoint, SFS data has the benefit that it can be obtained from pooled sequence data and does not incur the cost or possible introduction of error due to haplotype phasing (Liu and Fu, 2015; Anand et al., 2016). As a consequence, for many non-model species, SFS data will remain the most prevalent type (Han et al., 2013). In Simon and Huttley (2021) a method was given for reducing bias resulting from sequencing error in the calculation of the statistic *ρ* used for testing evolutionary neutrality. This involved modifying the likelihood equation to remove low frequency variants from the analysis. This approach could also be applied to the MCMC method described in this paper by modifying the likelihood equation (2) in the same way.

Bhaskar and Song (2014) have analysed an idealised situation in which the SFS data is produced by some specific defined stochastic generative process and the expected value of the SFS taken over multiple runs of this process is known with precision. They showed that any reasonable demographic history can be inferred from the expected value of the SFS under these conditions. On the other hand, the present paper deals with scenarios closer to the “real world” in which one only has knowledge of SFS data resulting from a single run of an unknown process. This necessarily results in uncertainty, which is expressed in the shape of the posterior distribution. Nevertheless, the results of Bhaskar and Song (2014) suggest that the SFS can be considered to approach sufficiency in some limiting sense.

### Summary and future work

We have proposed a method for computing posterior distributions for the coalescent times associated with a sample of sequences. The method does not make assumptions about the generative process for trees, but does assume that it is independent of mutation, that is, all mutations are selectively neutral. We used a Bayesian MCMC approach based on sampling from the space of TPCs, or tree matrices, jointly with relative branch lengths. This is effective because the likelihood function for the SFS can be computed directly from these parameters and sampling from them is equivalent to sampling from the full space of trees for the purpose of computing this likelihood. The relationship of the combinatorial structure of coalescent trees to the SFS was highlighted in Griffiths and Tavaré (1998). While that paper and the numerous other papers utilising its methods generally focused on computing expectations taken over the space of tree matrices using equation (3.3) of Griffiths and Tavaré (1998), we have taken a sampling approach which facilitated the use of Markov Chain Monte Carlo techniques. We also defined novel prior distributions for this purpose, covering the case where there is a belief as to the demographic history of the population that produced the data as well as the case where there is none.

A strength of the Bayesian method is that the joint distribution of branch lengths can be derived. This allows for representation of ancestral probabilities and other related quantities. The cost of this is a “curse of dimensionality” effect as the number of parameters to be estimated (the branch lengths) increases with sample size. This is intrinsic to the problem of estimating the full complement of branch lengths, rather than being specific to this method.

The use of Bayesian methods provides a clear indication of the degree of uncertainty in results arising from the stochastic nature of mutation and from our lack of knowledge of the actual tree structure. Both the simulation and Nuu-Chah-Nulth results demonstrate that the posterior distributions of branch lengths or coalescent times can be quite highly dispersed. It follows that point estimates should be treated with caution and credible intervals preferred.

There are a number of potential areas for future work extending the above methods. We have shown that Thomson’s estimator for the TMRCA does not suffer from the “curse of dimensionality” to the same degree as our Bayesian approach. It may be possible to combine the benefits of the two approaches, by using the Thomson estimate of the TMRCA to condition the prior distribution used in a Bayesian analysis. Another broad extension is to seek to estimate the demographic history of a population from the SFS. The simplest approach is to use the MMSE estimate of branch lengths in place of a maximum likelihood estimate in the calculation of skyline plots as presented in Pybus et al. (2000) and Strimmer and Pybus (2001).

## ACKNOWLEDGMENTS

This research was supported by an Australian Government Research Training Program (RTP) Scholarship to HS. The authors would like to thank John Wakeley for helpful comments.

## REFERENCES

Aitchison, J. (1982). The statistical analysis of compositional data. Journal of the Royal Statistical Society. Series B (Methodological), 44(2):139–177.

Al-Rfou, R., Alain, G., Almahairi, A., Angermueller, C., Bahdanau, D., Ballas, N., Bastien, F., Bayer, J., Belikov, A., Belopolsky, A., et al. (2016). Theano: A python framework for fast computation of mathematical expressions. arXiv e-prints, pages arXiv–1605.

Anand, S., Mangano, E., Barizzone, N., Bordoni, R., Sorosina, M., Clarelli, F., Corrado, L., Boneschi, F. M., D’Alfonso, S., and De Bellis, G. (2016). Next generation sequencing of pooled samples: guideline for variants’ filtering. Scientific Reports, 6(1):1–9.

Bhaskar, A. and Song, Y. S. (2014). Descartes’ rule of signs and the identifiability of population demographic models from genomic variation data. Annals of Statistics, 42(6):2469–2493.

Devroye, L. (1986). Non-uniform random variate generation. Springer-Verlag, New York.

Donnelly, P., Tavaré, S., Balding, D. J., and Griffiths, R. C. (1996). Estimating the age of the common ancestor of men from the ZFY intron. Science, 272(5266):1357–1359.

Dorit, R. L., Akashi, H., Gilbert, W., et al. (1995). Absence of polymorphism at the ZFY locus on the human Y chromosome. Science, 268(5214):1183–1185.

Dunson, D. B. and Tindall, K. R. (2000). Bayesian analysis of mutational spectra. Genetics, 156(3):1411–1418.

Ewens, W. J. (2004). Mathematical population genetics. I. Theoretical introduction. Number 27 in Interdisciplinary applied mathematics. Springer-Verlag, New York.

Felsenstein, J. (2005). Accuracy of coalescent likelihood estimates: do we need more sites, more sequences, or more loci? Molecular Biology and Evolution, 23(3):691–700.

Felsenstein, J., Kuhner, M. K., Yamato, J., and Beerli, P. (1999). Likelihoods on coalescents: a Monte Carlo sampling approach to inferring parameters from population samples of molecular data. Lecture Notes-Monograph Series, pages 163–185.

Gelman, A., Carlin, J. B., Stern, H. S., Dunson, D. B., Vehtari, A., and Rubin, D. B. (2014). Bayesian data analysis. CRC press Boca Raton, FL, 3rd edition.

Griffiths, R. and Tavaré, S. (1994). Ancestral inference in population genetics. Statistical Science, 9(3):307–319.

Griffiths, R. and Tavaré, S. (1998). The age of a mutation in a general coalescent tree. Stochastic Models, 14(1-2):273–295.

Gusfield, D. (1991). Efficient algorithms for inferring evolutionary trees. Networks, 21(1):19–28.

Han, E., Sinsheimer, J. S., and Novembre, J. (2013). Characterizing bias in population genetic inferences from low-coverage sequencing data. Molecular Biology and Evolution, 31(3):723–735.

Hein, J., Schierup, M., and Wiuf, C. (2005). Gene genealogies, variation and evolution: a primer in coalescent theory. Oxford University Press, Oxford UK.

Hodgkinson, A., Ladoukakis, E., and Eyre-Walker, A. (2009). Cryptic variation in the human mutation rate. PLoS Biology, 7:e1000027.

Hudson, R. R. (2002). Generating samples under a Wright-Fisher neutral model of genetic variation. Bioinformatics, 18(2):337–338.

Hudson, R. R. (2007). The variance of coalescent time estimates from DNA sequences. Journal of Molecular Evolution, 64(6):702–705.

Hunter, J. D. (2007). Matplotlib: A 2D graphics environment. Computing in Science & Engineering, 9(3):90–95.

Huttley, G. (2016). scitrack 0.1.1. [https://pypi.org/project/scitrack/0.1.1/].

Jaynes, E. T. (2003). Probability theory: The logic of science. Cambridge University Press, UK.

Kelleher, J., Etheridge, A. M., and McVean, G. (2016). Efficient coalescent simulation and genealogical analysis for large sample sizes. PLoS Computational Biology, 12(5):e1004842.

Knight, R., Maxwell, P., Birmingham, A., Carnes, J., Caporaso, J. G., Easton, B. C., Eaton, M., Hamady, M., Lindsay, H., Liu, Z., et al. (2007). Pycogent: a toolkit for making sense from sequence. Genome Biology, 8(8):1–16.

Kuhner, M. K., Yamato, J., and Felsenstein, J. (1995). Estimating effective population size and mutation rate from sequence data using Metropolis-Hastings sampling. Genetics, 140(4):1421–1430.

Kumar, R., Carroll, C., Hartikainen, A., and Martin, O. (2019). ArviZ a unified library for exploratory analysis of Bayesian models in Python. Journal of Open Source Software, 4(33):1143.

Lehmer, D. H. (1960). Teaching combinatorial tricks to a computer. In Proc. Sympos. Appl. Math. Combinatorial Analysis, volume 10, pages 179–193.

Liu, X. and Fu, Y.-X. (2015). Exploring population size changes using SNP frequency spectra. Nature Genetics, 47(5):555–559.

Marjoram, P. and Tavaré, S. (2006). Modern computational approaches for analysing molecular genetic variation data. Nature Reviews Genetics, 7(10):759–770.

Markovtsova, L., Marjoram, P., and Tavaré, S. (2000). The age of a unique event polymorphism. Genetics, 156(1):401–409.

McKinney, W. (2010). Data structures for statistical computing in Python. In van der Walt, S.and Millman, J., editors, Proceedings of the 9th Python in Science Conference, pages 51–56.

Meligkotsidou, L. and Fearnhead, P. (2005). Maximum-likelihood estimation of coalescence times in genealogical trees. Genetics, 171(4):2073–2084.

Nielsen, R., Korneliussen, T., Albrechtsen, A., Li, Y., and Wang, J. (2012). SNP calling, genotype calling, and sample allele frequency estimation from new-generation sequencing data. PLoS ONE, 7(7):e37558.

Pedregosa, F., Varoquaux, G., Gramfort, A., Michel, V., Thirion, B., Grisel, O., Blondel, M., Prettenhofer, P., Weiss, R., Dubourg, V., Vanderplas, J., Passos, A., Cournapeau, D., Brucher, M., Perrot, M., and Duchesnay, E. (2011). Scikit-learn: Machine learning in Python. Journal of Machine Learning Research, 12:2825–2830.

Peter, B. M., Huerta-Sanchez, E., and Nielsen, R. (2012). Distinguishing between selective sweeps from standing variation and from a de novo mutation. PLoS Genetics, 8(10):e1003011.

Przeworski, M. (2003). Estimating the time since the fixation of a beneficial allele. Genetics, 164(4):1667–1676.

Pybus, O. G., Rambaut, A., and Harvey, P. H. (2000). An integrated framework for the inference of viral population history from reconstructed genealogies. Genetics, 155(3):1429–1437.

Ronacher, A. (2009). click 7.0. [https://pypi.org/project/click/].

Rose, E. and Bayles, B. (2012). more-itertools 8.7.0. https://github.com/more-itertools/more-itertools.

Rudin, W. (1987). Real and complex analysis. McGraw-Hill, New York.

Salvatier, J., Wiecki, T. V., and Fonnesbeck, C. (2016). Probabilistic programming in Python using PyMC3. PeerJ Computer Science, 2:e55.

Simon, H. and Huttley, G. A. (2020). Quantifying influences on intragenomic mutation rate. G3: Genes, Genomes, Genetics, 10(8):2641–2652.

Simon, H. and Huttley, G. A. (2021). A new likelihood-based test for natural selection. bioRxiv.

Stephens, M. and Donnelly, P. (2000). Inference in molecular population genetics. Journal of the Royal Statistical Society: Series B (Statistical Methodology), 62(4):605–635.

Strimmer, K. and Pybus, O. G. (2001). Exploring the demographic history of DNA sequences using the generalized skyline plot. Molecular Biology and Evolution, 18(12):2298–2305.

Tajima, F. (1983). Evolutionary relationship of DNA sequences in finite populations. Genetics, 105(2):437–460.

Tang, H., Siegmund, D. O., Shen, P., Oefner, P. J., and Feldman, M. W. (2002). Frequentist estimation of coalescence times from nucleotide sequence data using a tree-based partition. Genetics, 161(1):447–459.

Tavaré, S. (1984). Line-of-descent and genealogical processes, and their applications in population genetics models. Theoretical Population Biology, 26(2):119–164.

Tavaré, S., Balding, D. J., Griffiths, R. C., and Donnelly, P. (1997). Inferring coalescence times from DNA sequence data. Genetics, 145(2):505–518.

Thomson, R., Pritchard Jonathan, K., Shen, P., Oefner Peter, J., and Feldman Marcus, W. (2000). Recent common ancestry of human Y chromosomes: Evidence from DNA sequence data. Proceedings of the National Academy of Sciences, 97(13):7360–7365.

Virtanen, P., Gommers, R., Oliphant, T. E., Haberland, M., Reddy, T., Cournapeau, D., Burovski, E., Peterson, P., Weckesser, W., Bright, J., et al. (2020). Scipy 1.0: fundamental algorithms for scientific computing in python. Nature Methods, 17(3):261–272.

Wakeley, J. (2009). Coalescent theory: an introduction. Roberts & Company, Green-wood Village Colorado.

Walsh, B. (2001). Estimating the time to the most recent common ancestor for the Y chromosome or mitochondrial DNA for a pair of individuals. Genetics, 158(2):897–912.

Ward, R. H., Frazier, B. L., Dew-Jager, K., and Pääbo, S. (1991). Extensive mitochondrial diversity within a single Amerindian tribe. Proceedings of the National Academy of Sciences, 88(19):8720–8724.

Waskom, M., Botvinnik, O., O’Kane, D., Hobson, P., Lukauskas, S., Gemperline, D. C., Augspurger, T., Halchenko, Y., Cole, J. B., Warmenhoven, J., de Ruiter, J., Pye, C., Hoyer, S., Vanderplas, J., Villalba, S., Kunter, G., Quintero, E., Bachant, P., Martin, M., Meyer, K., Miles, A., Ram, Y., Yarkoni, T., Williams, M. L., Evans, C., Fitzgerald, C., Brian Fonnesbeck, C., Lee, A., and Qalieh, A. (2017). Seaborn: v0.8.1. https://doi.org/10.5281/zenodo.883859.

Yang, Z., Li, J., Wiehe, T., and Li, H. (2018). Detecting recent positive selection with a single locus test bipartitioning the coalescent tree. Genetics, 208(2):791–805.

