## Supplementary Information for "Bayesian Inference of Joint Coalescence Times for Sampled Sequences"

#### 1 Demographic history used in simulation.

Demographic history for population simulation analysed in Figures 1 (c) and S3 (below).

```
================================
Epoch: 0 -- 10000.0 generations
================================
     start     end      growth_rate |     0
   -------- --------       -------- | --------
0 |  5e+04  2.49e+03         0.0003 |     0

Events @ generation 10000.0
- Population parameter change for -1: initial_size -> 30000
                                      growth_rate -> 0

==================================
Epoch: 10000.0 -- inf generations
==================================
     start     end      growth_rate |     0
   -------- --------       -------- | --------
0 |  3e+04    3e+04               0 |     0
```

#### 2 Example posterior distributions for sample size $n = 8$

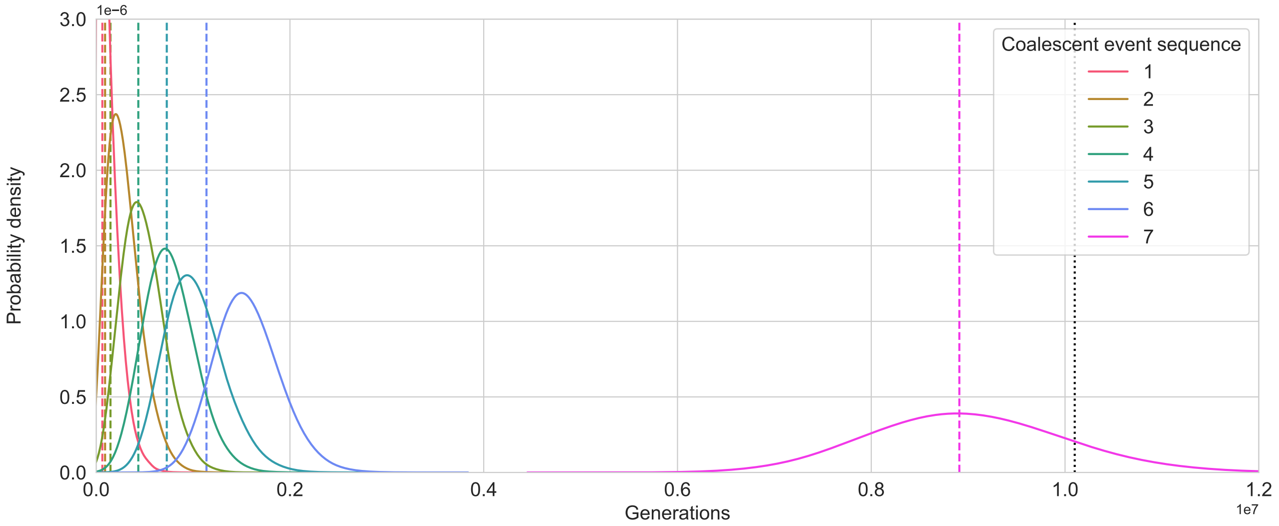


(a)

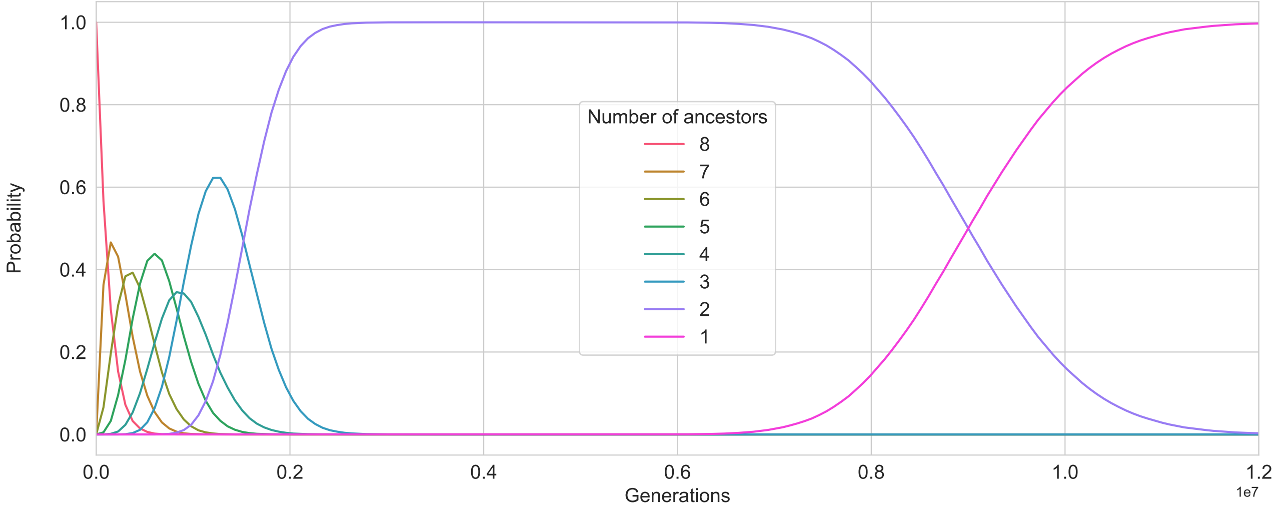


(b)

Figure S1: Two representations of a Bayesian analysis of a simulated sample (of size $n = 8$) using a constant size population model. The site frequency spectrum for the simulated sample is (52, 5, 5, 3, 0, 0, 45). (a) Posterior density functions of coalescence times. The numbers in the legend represent the order of coalescence times counting backward from the most recent coalescent event to the most distant (the TMRCA). The actual coalescence times being estimated, at which $k$ ancestors of the sample reduce to $k - 1$ ancestors, are shown as vertical lines. The Thomson estimator for the TMRCA is shown as a black dotted line. (b) The ancestral probability distributions. The plot for $k = 1, \ldots, 8$ (Legend) shows the posterior probability that there are exactly $k$ extant ancestors of the sample at the time $t$ shown on the horizontal axis.

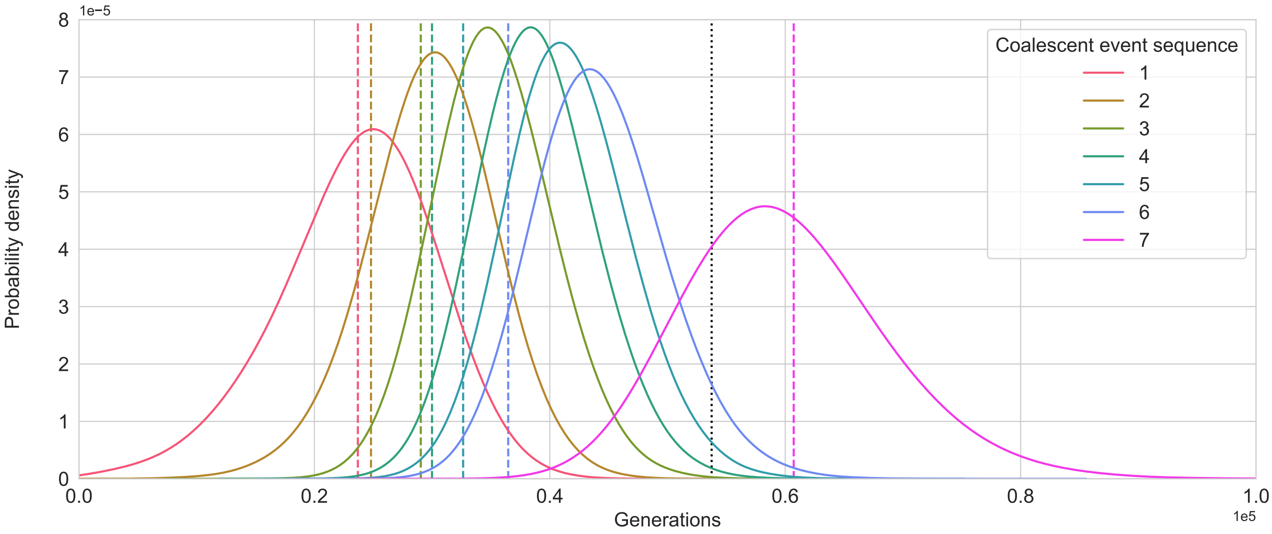


(a)

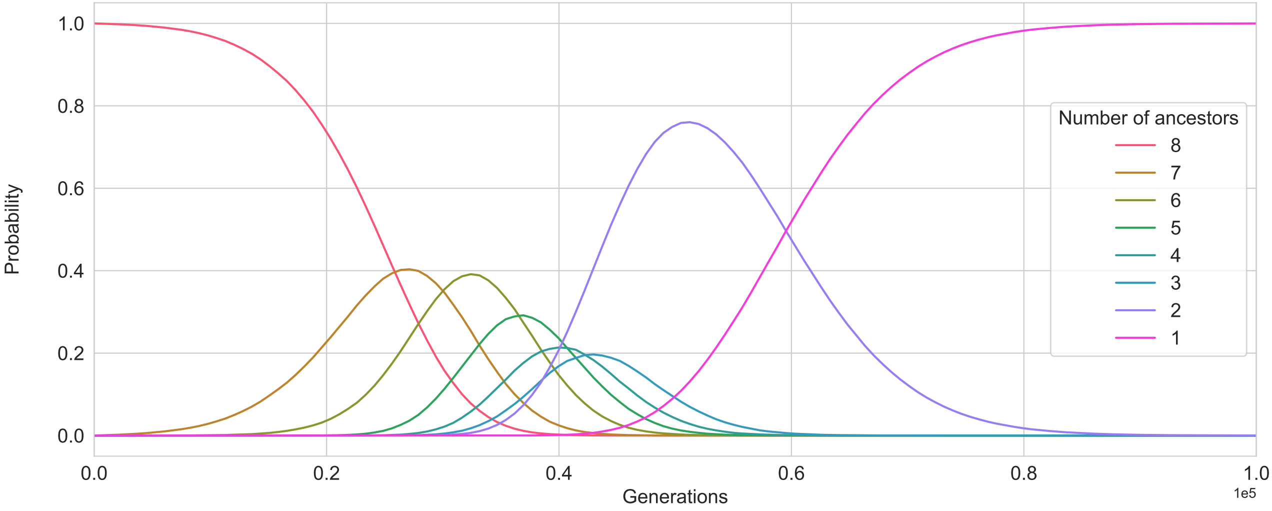


(b)

Figure S2: Two representations of a Bayesian analysis of a simulated sample (of size $n = 8$) using a a population model with growth parameter of 0.0001. The site frequency spectrum for the simulated sample is (91, 2, 2, 0, 0, 0, 4). (a) Posterior density functions of coalescence times. The numbers in the legend represent the order of coalescence times counting backward from the most recent coalescent event to the most distant (the TMRCA). The actual coalescence times being estimated, at which $k$ ancestors of the sample reduce to $k-1$ ancestors, are shown as vertical lines. The Thomson estimator for the TMRCA is shown as a black dotted line. (b) The ancestral probability distributions. The plot for $k = 1, \dots, 8$ (Legend) shows the posterior probability that there are exactly $k$ extant ancestors of the sample at the time $t$ shown on the horizontal axis

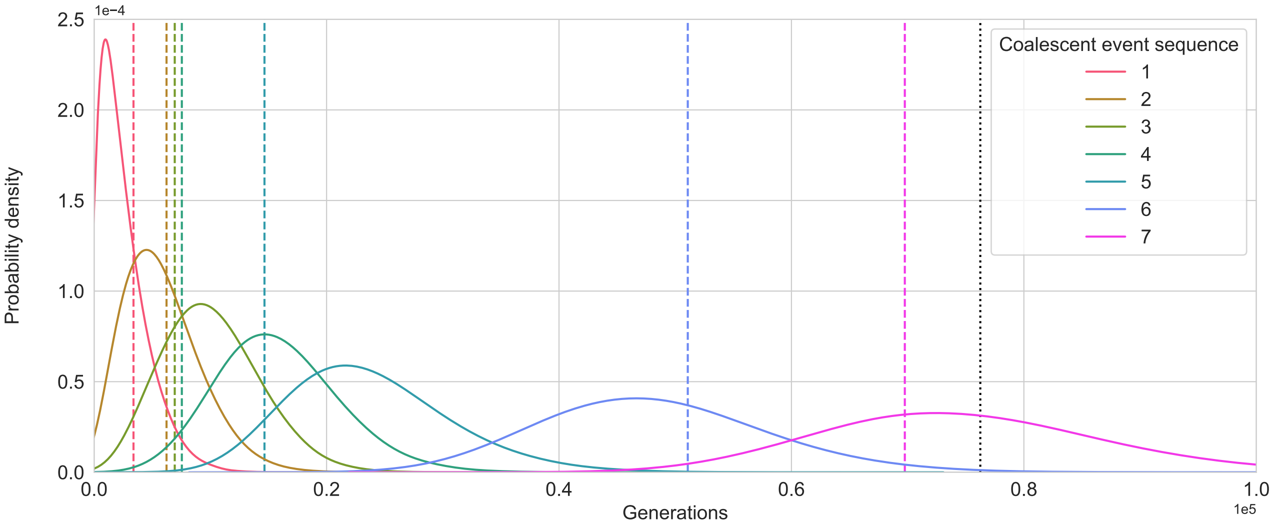


(a)

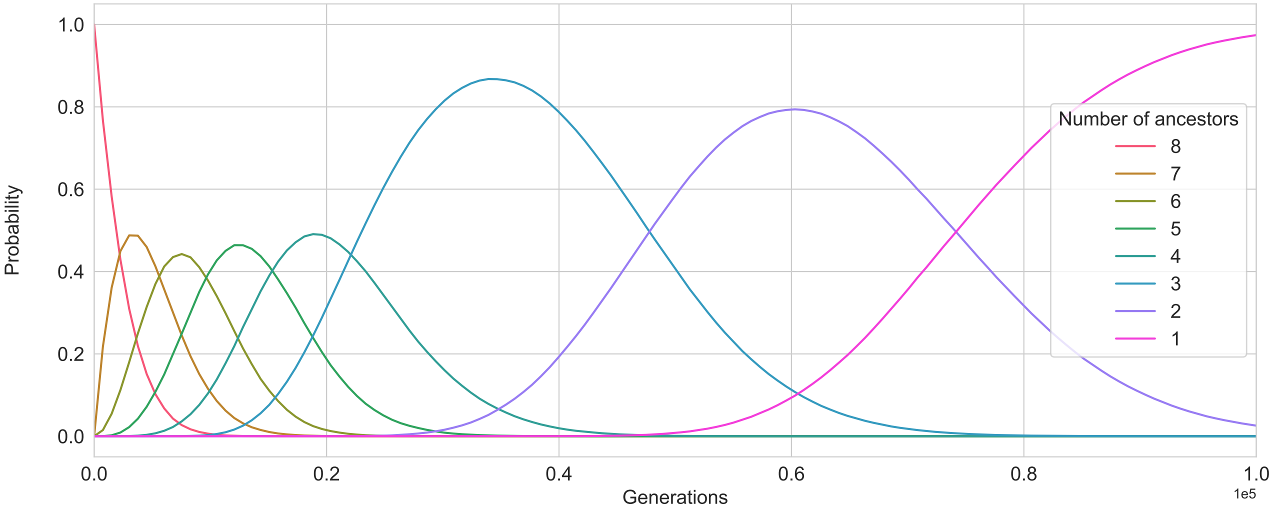


(b)

Figure S3: Two representations of a Bayesian analysis of a simulated sample (of size $n = 8$) using a demographic model involving a population bottleneck. (see Section 1 for details). The site frequency spectrum for the simulated sample is (20, 5, 13, 7, 5, 0, 0). (a) Posterior density functions of coalescence times. The numbers in the legend represent the order of coalescence times counting backward from the most recent coalescent event to the most distant (the TMRCA). The actual coalescence times being estimated, at which $k$ ancestors of the sample reduce to $k - 1$ ancestors, are shown as vertical lines. The Thomson estimator for the TMRCA is shown as a black dotted line. (b) The ancestral probability distributions. The plot for $k = 1, \ldots, 8$ (Legend) shows the posterior probability that there are exactly $k$ extant ancestors of the sample at the time $t$ shown on the horizontal axis

#### 3 Inference of basal split for Nuu-Chah-Nulth population sample

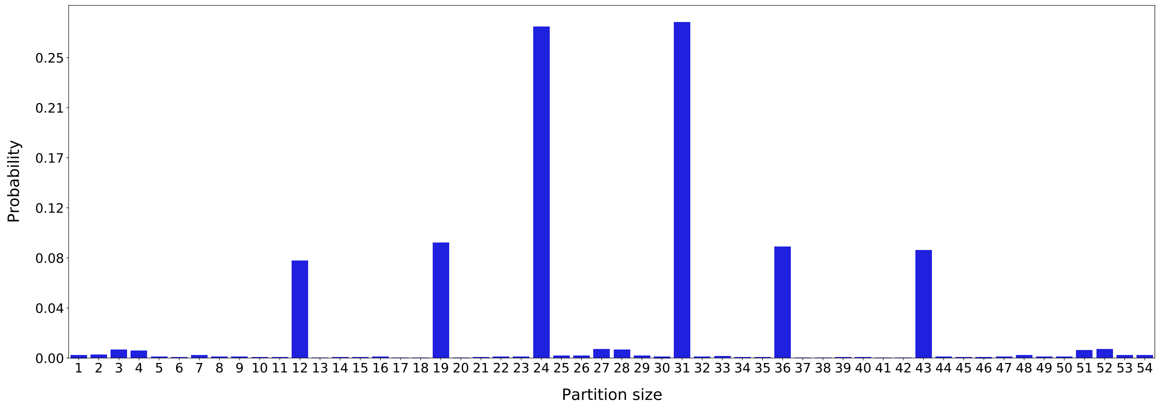


Figure S4: Posterior distributions of the basal split of coalescent trees inferred from Nuu-Chah-Nulth data. The y-axis shows the probability that a split occurred with the number of members of the sample to the left of the split being given by the x-value.

#### 4 Thomson's estimator derived by the method of moments

We show that Thomson's estimator (Thomson et al., 2000) can be derived using the method of moments from the same model used in our Bayesian inference approach.

Let $\mathbf{s} = (s_1, \ldots, s_{n-1})$ be the site frequency spectrum for a sample of size $n$, i.e. $s_i$ is the number of mutations occurring $i$ times in the sample. Define $s_{k,j}$ as the number of mutations occurring $j$ times in the sample that occurred in coalescence period $2 \leq k \leq n$. Then:

$$s_j = \sum_{k=2}^{n} s_{k,j}$$

Define $d$ by:

$$d = \sum_{j=1}^{n-1} j s_j = \sum_{j=1}^{n-1} j \sum_{k=2}^{n} s_{k,j}. \tag{1}$$

Let $\psi$ be the unknown tree that generated the sample. Then, in our model $s_{k,j}$ is considered to be generated by the Poisson distribution $\text{Poi}(\mu t_k A_\psi[k-1, j])$, since $A_\psi[k-1, j]$ is defined as the number of branches at level $k$ with $j$ descendants. Hence if, substituting for $s_{k,j}$ in equation(1), we define:

$$D \sim \sum_{j=1}^{n-1} j \sum_{k=2}^{n} \text{Poi}(\mu t_k A_\psi[k-1, j]); \text{ we have:}$$

$$\mathbb{E}(D) = \sum_{j=1}^{n-1} j \sum_{k=2}^{n} \mu t_k A_\psi[k-1, j] = \mu \sum_{k=2}^{n} t_k \sum_{j=1}^{n-1} j A_\psi[k-1, j] = \mu \sum_{k=2}^{n} t_k n$$

where the last equality results from equation (1) of Simon and Huttley (2021). Since we also have $\mathbb{E}(D) = d$, we obtain the estimate:

$$\widehat{\sum_{k=2}^{n} t_k} = \frac{d}{n\mu} = \frac{\sum_{j=1}^{n-1} j s_j}{n\mu}$$

### References

Simon, H. and Huttley, G. A. (2021). A new likelihood-based test for natural selection. *bioRxiv*.

Thomson, R., Pritchard, Jonathan, K., Shen, P., Oefner, Peter, J., and Feldman, Marcus, W. (2000). Recent common ancestry of human Y chromosomes: Evidence from DNA sequence data. *Proceedings of the National Academy of Sciences*, 97(13):7360–7365.
